# Vocal species are more central in Eastern Himalayan Mixed-Species bird flocks

**DOI:** 10.1101/2024.12.06.626929

**Authors:** Satyam Gupta, Akshay Bharadwaj, Akshiti Bhat, Aiti Thapa, Aman Biswakarma, Bharat Tamang, Binod Munda, Biren Biswakarma, Dema Tamang, Dambar Pradhan, Mangal K Rai, Rohit Rai, Shambhu Rai, Umesh Srinivasan

**Affiliations:** Department of Ecology and Environmental Sciences, Pondicherry University, India; Conservation Biology Lab, University of Neuchâtel, Switzerland; Organismic and Evolutionary Biology Lab, University of Massachusetts Amherst, USA; Centre for Ecological Sciences, Indian Institute of Science, Bangalore, India

**Keywords:** Animal Communication, Bioacoustics, Eastern Himalaya, Mixed-species Flock, Network Analysis, Social Organization

## Abstract

Mixed-species flocks (MSF) represent an important form of social organisation in bird communities worldwide. Despite its likely importance in flock formation and cohesion, the role of vocal communication in the formation and maintenance of MSF in birds is hitherto understudied. In this study, we examine if a species’ centrality within a mixed-species flock is influenced by its vocal behaviour during the dawn chorus, i.e., the time of MSF formation. Using acoustic sampling and field observations, we studied the bird species found in MSF in the Eastern Himalayas. Our results show differential vocal activity patterns among MSF-forming bird species and suggest a positive correlation between calling rates and closeness centrality (species’ importance in a flock) in understory MSFs. We also found a more synchronised vocalisation pattern in the understory MSFs, with a consistent peak in vocal activity in the early morning hours, whereas no consistent vocal pattern was found for canopy flocks. Overall, our results suggest a potential mechanism that drives MSF formation wherein the vocal activity of central species precedes and likely attracts participation from other attendant species.

## INTRODUCTION

Multi-species associations are common in nature. Interspecific interactions consisting of individuals of two or more species can aid in predator avoidance and enhance foraging success (Morse 1977; Sridhar et al. 2009; Goodale et al. 2017). Mixed-species flocks (hereafter, MSF or “flocks”) are interspecific associations of two or more species, representing an important feature of bird communities worldwide (Greenberg 2000; Harrison and Whitehouse 2011).

MSF consist of species exhibiting a continuum of social cohesion within flocks, ranging from loose aggregations such as those of herons and seabirds (Sealy 1973, Baltz 1977, Kushlan 1977) to highly structured winter foraging groups of chickadees (*Parus sp.*) (Hogstad 1978) and multi-species territoriality in the Amazon (Jullien and Thiollay 1998). The formation and maintenance of these flocks are influenced by a complex interplay of factors, including species-specific ecological traits, foraging strategies, and social behaviours. While traditional models categorized species into rigid roles such as ’nuclear’ and ’satellite’ (Winterbottom 1943; Powell 1985; Hutto 1994; Farley et al. 2008), more recent research has highlighted the dynamic nature of these roles and the potential for significant behavioural flexibility in response to changing environmental conditions and social interactions (Dolby & Grubb, 2000; Srinivasan et al. 2010, Borah et al. 2018).

While several studies have explored the composition of MSF, the vocal behaviour of its constituent species remains understudied. Acoustic communication plays a pivotal role in forming and maintaining MSF (Suzuki 2012). Bird vocalisations can serve multiple functions, ranging from species recognition to maintenance of interactions and alarm calling (Marler, 2004). Indeed, some bird species have been shown to produce specific calls to attract others to join MSF (Diamond 1987, Goodale & Kotagama, 2006). In the context of MSF, these distinct and recognisable calls can allow individuals to locate and join the flock. By investigating the patterns in species-specific vocalizations, we can elucidate how these signals contribute to flock formation, cohesion and information exchange (Goodale & Kotagama, 2005).

In this study, we examined the role of vocalisation patterns in forming and maintaining MSF. By examining temporal patterns in species-specific vocalizations, we aim to understand how vocalisation could influence the role of each species in the MSF formation process. Our *a priori* hypothesis was that species exhibiting higher calling rates would be more central in the MSF networks than species with lower calling rates.

## MATERIALS AND METHODS

### STUDY AREA

We conducted this study in the Eaglenest Wildlife Sanctuary, located in the West Kameng district of Arunachal Pradesh, India (27.07°N; 92.40°E), which is situated within the Eastern Himalaya global biodiversity hotspot (Figure 1). We sampled MSF in the montane wet temperate forest, ∼2000 m above sea level. The forest is characterised by dominant tree species such as *Quercus lamellosa*, *Alnus nipalensis*, and *Michelia doltsopa* in the canopy, along with an understorey dominated by bamboo species (*Chimonobambusa sp.*; Srinivasan, 2019).

**Figure 1:**
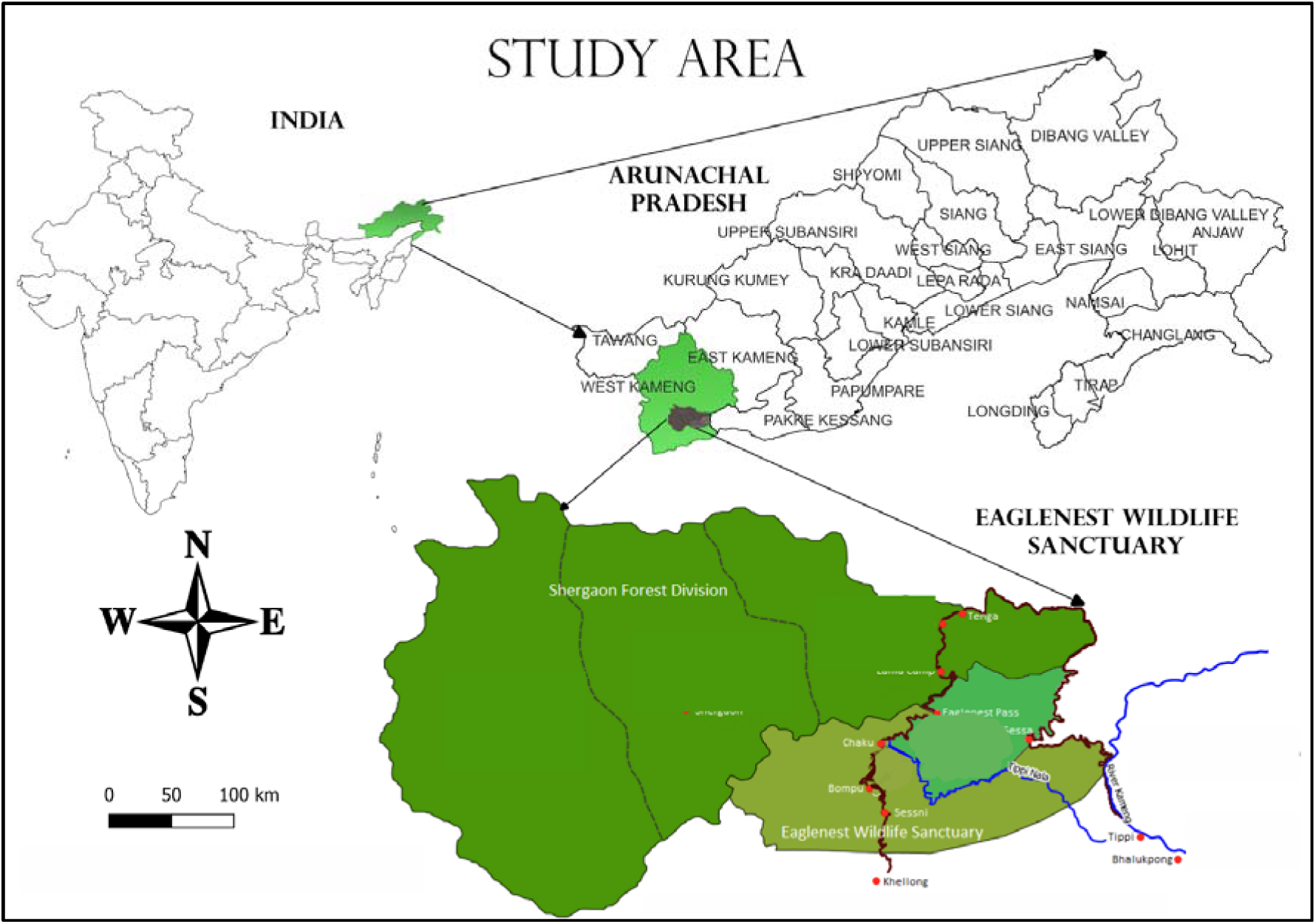
A map of the study area showing the location of Eaglenest Wildlife Sanctuary, India.

### ETHICAL NOTE

All procedures complied with the ASAB/ABS principles. Our study was confined to noninvasive and non-disruptive behavioral observation techniques. Given the non-invasive nature of our research, no animals were harmed, and it did not require review from an institutional or governmental regulatory body. The Arunachal Forest Department granted permission for the study under permit number CWLW/GEN/998/2023/Pt-VIII(B)/3493-96.

### SAMPLING METHODS

We used two sampling methods for data collection:

#### 1. Mixed-Species Flock Observation

We observed MSF between 15^th^ January and 11^th^ March 2024 at Eaglenest Wildlife Sanctuary near Bompu camp. An MSF is a single observation of a group of different species of birds moving together over space and time (usually at least 5 minutes). We focused on two stretches of a fairweather road, where observers monitored MSF daily (except during rain) from 6:00 AM to 10:00 AM and 2:00 PM to 4:00 PM.

Mixed-species flocks were defined as groups of birds from different species foraging and moving together within a short distance (<=5m) of each other. For each flock encountered, we recorded the number of species, the number of individuals of each species, the flock type (based on their habitat stratum), time, and location (recorded using a GARMIN Etrex 30x GPS device) for each observed flock. We used a Sony ICD-PX240 MP3 recorder for data collection, which proved more efficient than manual recording due to the dynamic nature of MSF. Our observations were confined to road stretches for improved species identification. To enhance the network analysis, we incorporated MSF data collected in 2023 at the same site and during the same season, with the 2024 data.

#### 2. Acoustic Sampling

##### 2.1. Active Acoustic Sampling

We collected “active” acoustic data from mixed-species flocks passing through specific locations along the stretch of Fairweather Road (detailed in Figure 2). Acoustic sampling was done between January 15th and March 11th, 2024. Active sampling involved us being present and adjusting the microphone’s orientation to optimize recording as the flocks moved (Britzke 2002). Our sampling took place from 6:00 AM to 10:00 AM each day. We had two teams stationed on different roads near Bompu camp, using Audiomoth recorders (Hill et al., 2019) for active acoustic sampling. At the same time as recording, observers documented flock composition, time, and GPS location using a custom Epicollect5 form (Aanensen et al., 2009) (Eaglenest Acoustic Sampling SAS).

**Figure 2.**
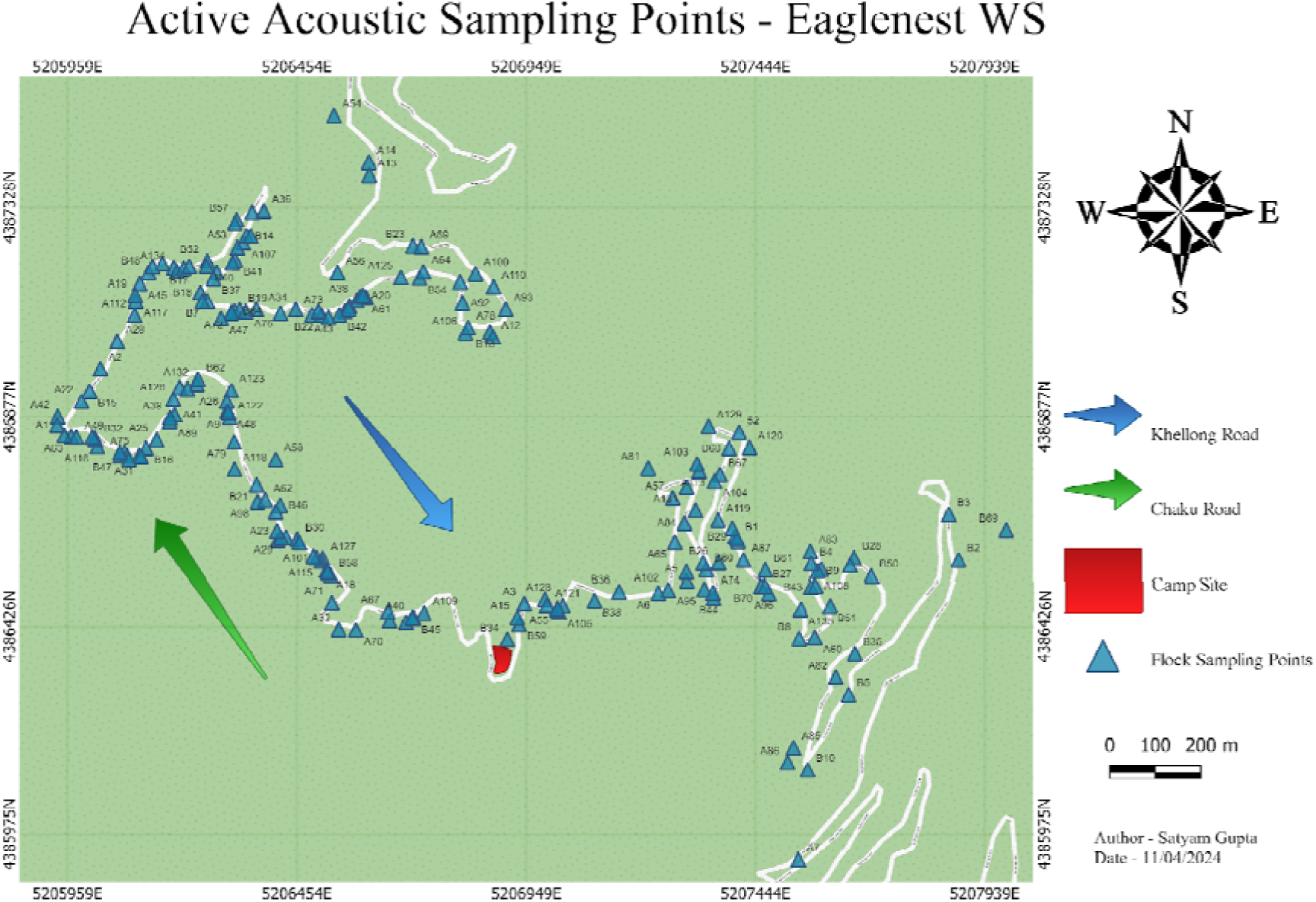
Active Acoustic Flock Sampling Locations

##### 2.2 Opportunistic Acoustic Sampling

We collected additional acoustic recordings opportunistically from MSF-participating bird species by recording individual calling bouts from January 15th to March 15th, 2024. We conducted this sampling daily from 6:00 AM to 10:00 AM Indian Standard Time (UTC +5:30), alongside the mixed-species flock observation detailed above.

#### 3. Preliminary Results

We observed a total of 188 MSF during the study period, comprising 55 unique species. This observational data was combined with the 2023 dataset of 280 MSF collected in the same study site and methodology. In total, we had species-level flock composition data for 347 MSFs (280 in 2023 and 188 in 2024) comprising 68 unique species. In the active acoustic sampling, we observed a total of 204 MSFs, comprising 40 unique species. Additionally, we opportunistically collected 1804 unique individual vocalization bouts of birds participating in MSFs.

### ANALYSIS

#### 1. Social Network Analysis

The MSF data were filtered to remove species encountered in less than ten observed flocks, following recommendations from Farine & Whitehead (2015). We used a walk-run clustering algorithm, implemented using the ’cluster’ package (v1.3.8; Maechler et al., 2023) in R statistical software (v4.3.2; R Core Team, 2023), which facilitates the classification of flocks into distinct clusters based on species co-occurrences. The ’igraph’ package (v2.1.3; Csardi & Nepusz, 2006) in R statistical software (v4.3.2; R Core Team, 2023) was used to construct species-level networks, where species were represented as nodes and co-occurrences as edges. Additionally, the ’igraph’ package (v2.1.3; Csardi & Nepusz, 2006) was also employed to calculate the closeness centrality of each species in the MSF data.

We have used closeness centrality to evaluate the relative position of species in flock networks, due to its suitability in traditional information-sharing property (Wey & Blumstein, 2010). The closeness centrality of a node (here, species) in a graph is the inverse of the average shortest path distance from the node to other nodes in the graph (Wasserman & Faust, 1994). It can be viewed as the efficiency of each node (species) in spreading information to all other nodes (species) (Wasserman & Faust, 1994). The higher the closeness centrality of a node, the shorter the average distance from the node to any other node, and thus the better positioned the node is in potentially spreading information to other nodes (Wasserman & Faust, 1994). All analysis was done in the R statistical software (v4.3.2; R Core Team 2023).

#### 2. Acoustic Analysis

Each MSF recording was segmented into one-minute files, and MSF species vocalizing within these segments were manually annotated through visual inspection of spectrograms and audio analysis, with corresponding marked timestamps. This process was systematically repeated for all the observed MSF. Acoustic analysis was done using Raven Pro (v1.6.1; Charif et al., 2010).

By combining opportunistic acoustic data with active sampling data, we were able to gather a more comprehensive dataset for calculating the various vocal parameters of a species. This integrated dataset included the species name, date, and time of vocalization for each recording. We organised the vocalization data for each species into 30-minute intervals, creating eight-time windows between 6:00 AM and 10:00 AM (6:00 - 6:30 A.M. as Window#1). Subsequently, we calculated the number of calls in all windows and identified the time window with the highest number of call counts as the “peak vocalization period” or “peak vocalization category”. We used the ’dplyr’ package in R Statistical Software (v4.3.2; R Core Team 2023) for data manipulation. We used the ’ggplot2’ package (Wickham et al., 2016) in R Statistical Software (v4.3.2; R Core Team 2023) to create heatmaps for visualising vocal activity patterns.

#### 3. Correlation Analysis

We analyzed the correlation between acoustic metrics and closeness centrality in two flock types out of three: the canopy flock (12 species analyzed) and the understory flock (9 species). We didn’t include the mid-story flock in the analysis since it consisted of only two species overall. The flock types were defined based on the social network analysis and aligned with prior natural history observations at the site.

We performed Pearson correlation tests in R Statistical Software (v4.3.2; R Core Team 2023) to examine relationships between social network properties (namely, closeness centrality of a species) and the vocal activity patterns (peak vocalisation period and mean call count). We visualised the results using the ’ggplot2’ package (Wickham et al., 2016).

### RESULTS

#### 1. Social Network Analysis

We analysed mixed-species flock data across 35 species observed during our data collection. We generated a matrix showing which species tend to co-occur and used this information to build the network at the species level.

Subsequently, using the network clustering algorithm, we identified three distinct types of clusters (Figure 3), which corresponded to the MSFs as follows -

1. Understory Flock (12 Species): **Understory flocks** consisted of flocks led by the Yellow-throated Fulvetta (*Schoeniparus cinereus*) observed during field observation. Species belonging to this flocktype are the Yellow-throated Fulvetta (*Schoeniparus cinereus*), Rufous-capped Babbler (*Cyanoderma ruficeps*), Golden Babbler (*Cyanoderma chrysaeum*), Grey-cheeked Warbler (*Phylloscopus poliogenys*) and other species. The full list of species can be found in Table 1 (Supplementary Data).
2. Midstory Flock – 3 Species **Mid-story flocks** were led by the Rusty-fronted Barwing (*Actinodura egertoni*) observed during field observation. Species belonging to this flock type are Rusty-fronted Barwing (*Actinodura egertoni*), Coral-billed Scimitar Babbler (*Pomatorhinus ferruginosus*) and White-breasted Parrotbill (*Psittiparus ruficeps*).
3. Canopy Flock – 20 Species **Canopy flocks** were the most species-rich flock types found in the study area, with several flocks sometimes exceeding 20 species in number. They consisted of flocks led by Yellow-cheeked Tit (*Parus spilonotus*) and Yellow-browed Tit (*Sylviparus modestus*), as observed during field observation. Species belonging to this flock type include the Lesser Racket-tailed Drongo (*Dicrurus remifer*), Red-tailed Minla (*Minla ignotincta*), Ashy-throated Warbler (*Phylloscopus maculipennis*), Himalayan Cutia (*Cutia nipalensis*) and others species. The full list of species can be found in Table 1 (Supplementary Data).

**Figure 3:**
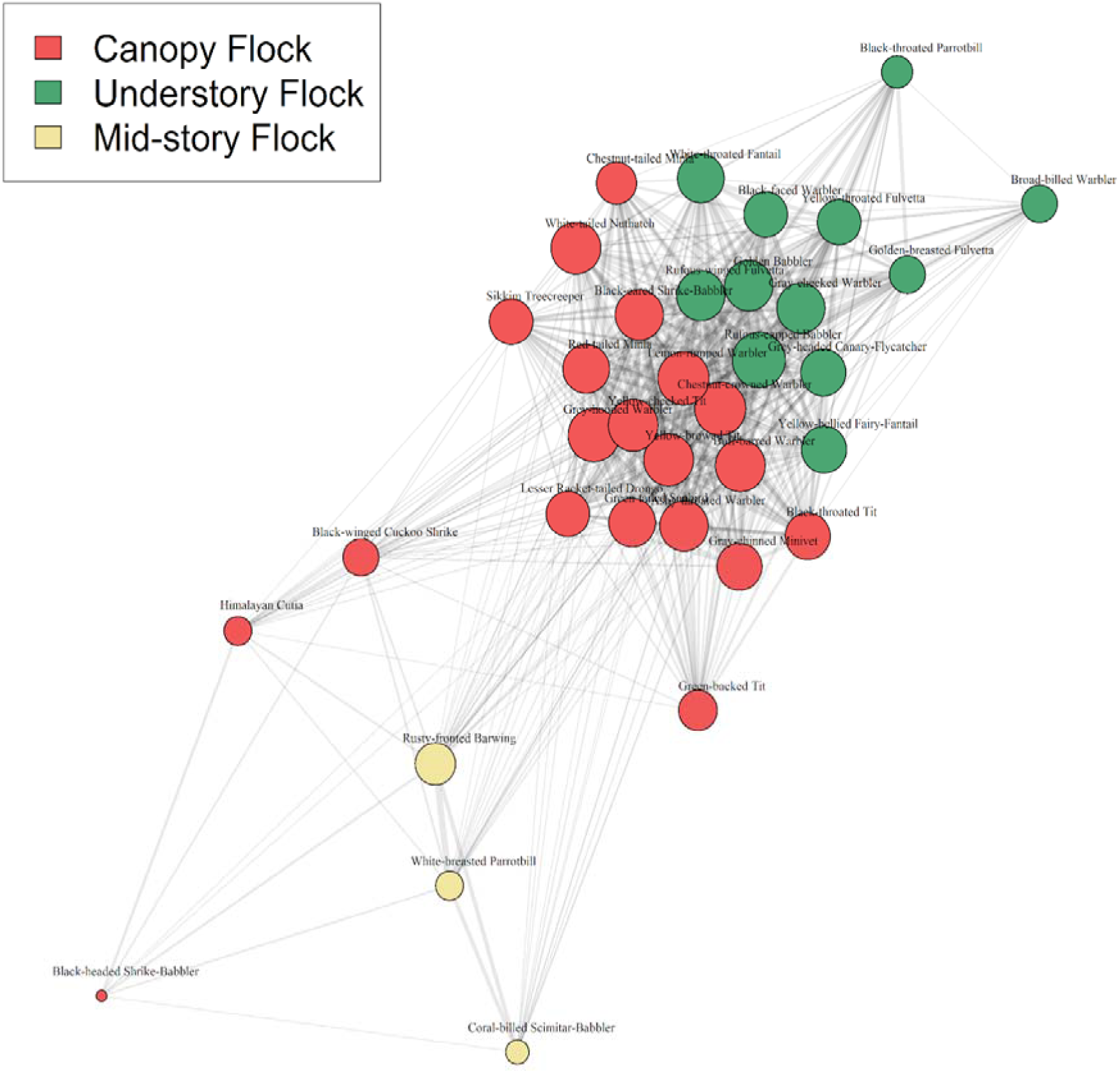
Species-level network visualising the co-occurrence of various species seen within mixed-species flocks. Each node represents a distinct bird species, and the edges (connected lines) between them represent relationships or interactions. The thickness of each edge represents the frequency of co-occurrence – a thicker edge signifies two species which are more commonly observed together. The network contains visible clusters of species (each “flock type”), and also shows strong connections between different types of flocks, indicating that while species may favour specific flock types (clusters), there are interactions across flock types.

#### 2. Vocal Activity Pattern

The acoustic analysis revealed distinct vocal activity patterns among species within the canopy and understory flocks (Figure 4). In the canopy flocks, species such as the Chestnut-crowned Warbler and Green-tailed Sunbird showed peak calling activity between 6:30 A.M. and 7:00 A.M. Other species, including the Ashy-throated Warbler and Himalayan Cutia, displayed moderate call activity across time, indicating a broader distribution of vocalization. In contrast, species such as the Lesser Racket-tailed Drongo (*Dicrurus remifer*) and Black-throated Tit (*Aegithalos concinnus*) vocalised less. These observations indicate that the canopy flock contains species with more diverse vocalization patterns compared with the understory flock, with some species being more active at specific times while others remain relatively inactive.

**Figure 4:**
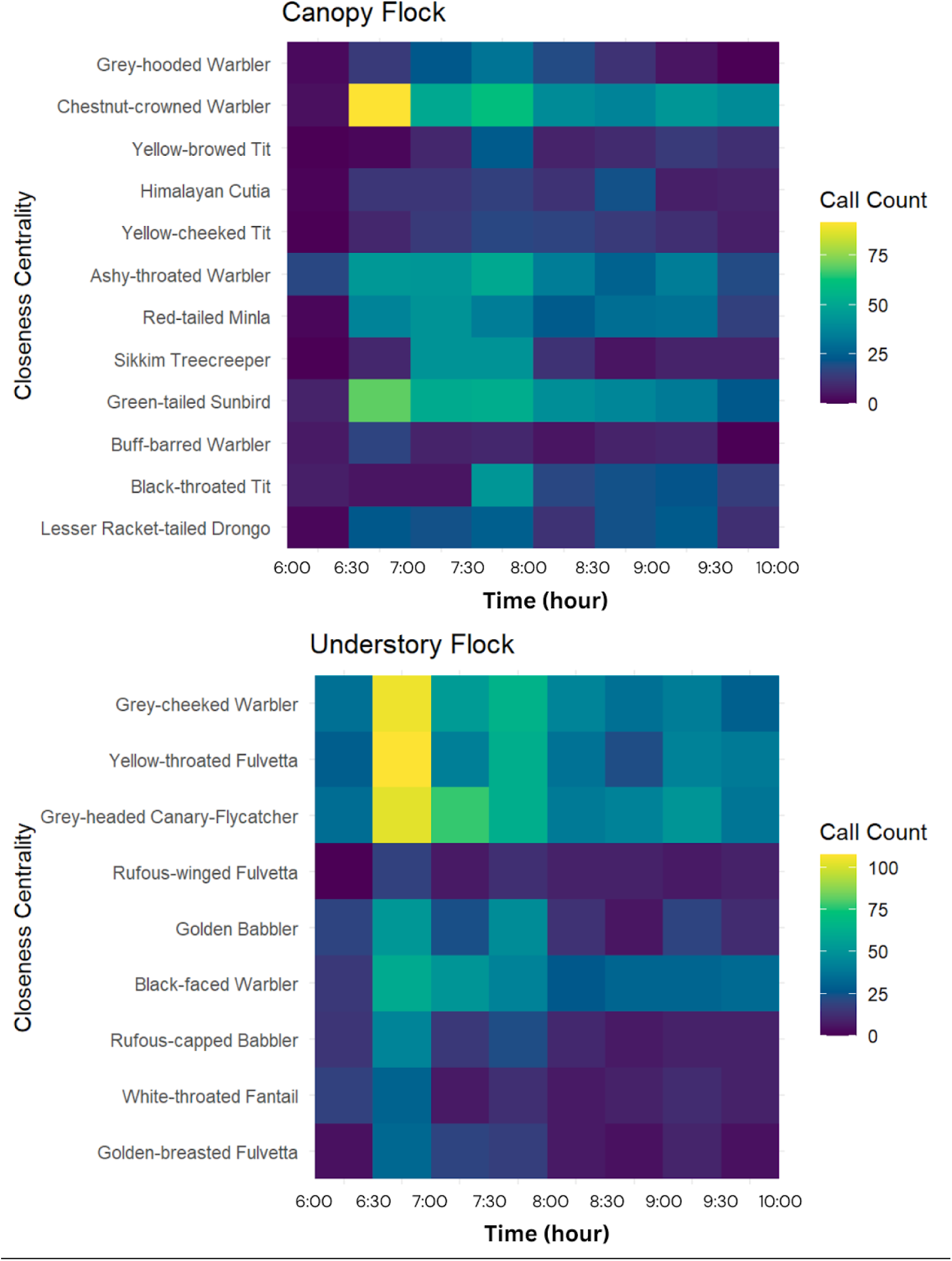
Vocal Activity Pattern: The heatmap displays the vocal activity patterns of different bird species in the canopy and understory flock types. The Y-axis represents species ordered in increasing closeness centrality values. The X-axis represents time in 30-minute intervals from 6:00 A.M. to 10:00 A.M. Each row corresponds to a specific bird species, and each column represents a 30-minute time interval. The colour intensity within each cell corresponds to the number of calls observed for that species during a specific vocal period.

In contrast, understory MSF birds exhibited earlier peak vocal activity as compared to the canopy MSF birds, with all the members vocalising highest between 6:30 A.M. and 7:00 A.M. The Yellow-throated Fulvetta and Grey-cheeked Warbler had particularly high call counts during this period, while species like the Grey-headed Canary-Flycatcher (*Culicicapa ceylonensis*) and Golden Babbler maintained moderate and consistent vocal activity throughout the day. Species such as the Golden-breasted Fulvetta and White-throated Fantail vocalised less. The understory flock generally followed a pattern of heightened morning activity, with different species showing peak calling activity as compared to the canopy flock.

#### 5. Correlation Analysis

Our analysis revealed different results for each type of MSF.

##### ***5.1*** Canopy Flock

Our analyses revealed a weak association between the peak vocalisation period, mean call count and closeness centrality. Correlation results indicated that the peak vocalization period explained a negligible amount of variance in closeness centrality (R² = 0.07, p = 0.62, F-values =0.14, df = 10). Similarly, mean call count exhibited a minimal relationship with closeness centrality (R² = 0.08, p = 0.79, F-values =0.06, df = 7).

##### ***5.2*** Understory Flock

In contrast, our analysis revealed a positive correlation between the mean call count and the closeness centrality of species within the understory flocks. (R² = 0.73; F-statistic = 8.10; p = 0.02) (Figure 5). However, the peak vocalization period was not tractable to statistical analysis due to its homogeneity within the understory flock. This lack of variability precluded its inclusion in the linear regression model. Furthermore, visual inspection (Suppl. Figure 4) revealed no discernible relationship between peak vocalization period and closeness centrality, suggesting that changes in the latter did not influence the former.

**Figure 5:**
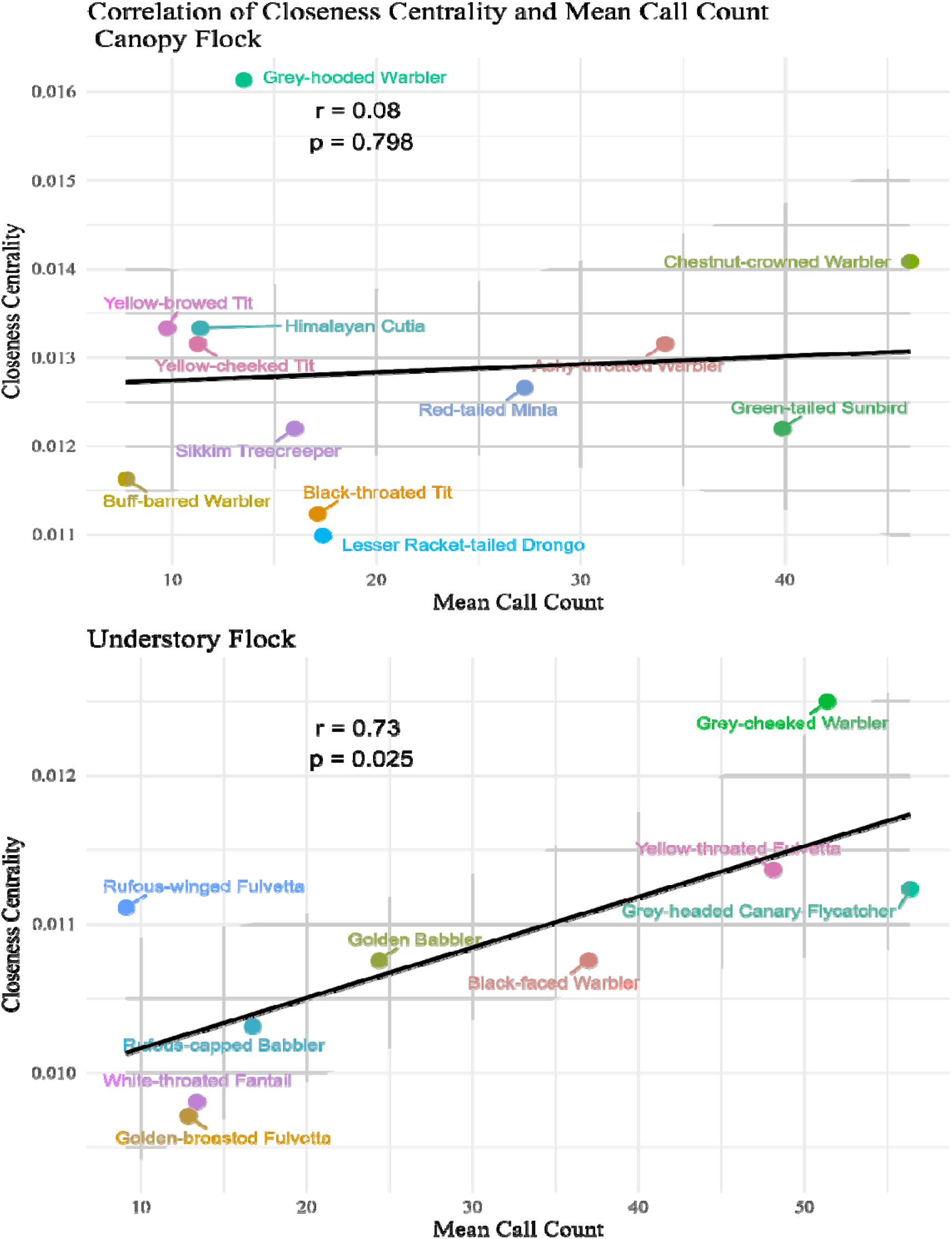
Correlation of Mean Call Count and Closeness Centrality. The Y-axis illustrates the values of closeness centrality, whereas the X-axis displays the mean call count over time. The plots indicate a positive correlation between closeness centrality and mean call count for understory flocks, suggesting that species with higher call counts are typically more central in understory flocks. This correlation is more pronounced in the Understory Flock, while it does not achieve statistical significance in the Canopy Flock.

## DISCUSSION

The phenomenon of MSF in birds has long fascinated researchers (Bates 1859); however, the mechanisms underpinning their formation and dynamics remain to be fully understood. Communication between different species plays a pivotal role in orchestrating the assembly and maintenance of these diverse avian congregations (Goodale et al., 2010).

Early research conducted by Munn (1985) and McDonald and Henderson (1977) provided insights into the significance of vocalization in the formation of MSF. Their findings indicated that certain bird species, such as titmice and antshrikes, produce loud calls in the morning, serving as a signal to attract other species to gather and form a flock. Once assembled, the vocalisation of the central species, often referred to as “leader species,” seems to reinforce flock cohesion and momentum (Bell, 1983; Diamond, 1987). These leaders usually initiate vocalizations that act as cues for others to follow, thereby influencing the structure and dynamics of the flock.

In addition, McDonald (1989) showcased the effectiveness of playback experiments and vocal muting techniques in gaining insight into the role of vocalization in flock formation. Monkkonen et al. (1996) undertook a thorough investigation on the impact of the territorial song of the willow tit (*Parus montanus*) on the creation of loose mixed feeding groups during the breeding season in Finland. Their observations of foraging aggregations among various species of boreal forest birds revealed varying responses to the willow tit’s song (Greenberg, 2000). They discovered that other boreal forest insectivores notably joined the willow tit in these mixed feeding groups (Greenberg, 2000). However, the level of distinction in the effect playback method was somewhat obscured due to experimental comparisons made with the song of a North American tyrant flycatcher and a piece of classical music (Greenberg, 2000). As a result, this study did not definitively uphold the idea that specific vocalizations are essential for promoting the formation of tit flocks (Greenberg, 2000).

Our research aimed to explore the connection between a species’ vocal behaviour during the crucial phase of flock formation and its importance within a flock.Overall, we identified three distinct types of flocks which are understory, canopy and midstory flocks in our study area, consistent with previous findings (Bharadwaj et al., 2023). Using network analyses, we identified several species, including the Yellow-bellied Fairy-fantail (*Chelidorhynx hypoxanthus*), Ashy-throated Warbler (*Phylloscopus maculipennis*), Grey-cheeked Warbler (*Phylloscopus poliogenys*), and White-throated Fantail (*Rhipidura albicollis*), that actively participated in multiple flock types, indicating high interspecies interaction and potential information sharing across different flock types. This dynamic interplay highlights the complexity of avian social networks and emphasizes the importance of considering individual species’ roles within larger ecological communities.

We postulated that species having higher vocalization rates, particularly during the early hours of the day when flock formation occurs are more important, or “central species”. Our hypothesis was based on the concept that central species play a crucial role in initiating and maintaining group cohesion through vocal communication. To investigate this hypothesis, we conducted field observations and acoustic recordings, meticulously documenting the vocal behaviour of different bird species within mixed-species flocks and their social interactions.

Our findings revealed intriguing patterns in vocal activity among different flock types. Birds in understory flocks, led by species such as the Yellow-throated Fulvetta (*Schoeniparus cinereus*), exhibited synchronized vocalization patterns, with a consistent peak in vocal activity during the early hours. However, we did not observe a clear pattern in canopy flock types, suggesting potential variation in the role of vocalizations in influencing MSF formation and cohesion.

Our study found that within the understory flock type, species with higher closeness centrality (referred to as central species) exhibited higher mean call counts, signalling that central species are more vocal than other flock members. These findings align with previous research, such as Smith (1991), which noted that key species such as tits (*Parus*) consistently vocalized within the flock. Similarly, in a tropical study, Wiley (1971) observed the dot-winged antwren, a central species in Central American antwren flocks, continuously vocalizing while moving between trees. This behaviour was used by other species to locate flocks as they moved slowly through dense vegetation, maintaining flock cohesion.

Additionally, Munn (1985) in the Neotropical forests of Peru, discovered that the leader species, the White-winged Shrike-tanager (*Lanio versicolor*), made chattering vocalizations, prompting other flock members to join in. Therefore, the higher calling rate of central species observed in our study may serve as a mechanism for maintaining the cohesion of other species and forming the flock. Our results highlight the crucial role of key species in initiating and maintaining flock cohesion through vocal communication.

Understanding the extent to which species exchange information with each other is crucial for comprehending the structure of animal groups. Measuring the level of information flow between species can offer valuable insights into the dynamics of species interactions that underpin group dynamics and may also indicate the extent to which certain species play a disproportionate role in maintaining group coherence.

In our research, while our primary focus was on the vocalization patterns and their correlation with the significance of species in mixed-species foraging (MSF) flocks, integrating metrics of information exchange between species could add further depth to our analysis. By quantifying the information exchange both within and between species, we can gain a better understanding of the communication dynamics within MSF, which in turn can provide valuable insights into the processes driving the formation and stability of these flocks.

Additionally, in terms of conservation, evaluating the symmetry of information flow can aid in identifying species that have key roles in maintaining group cohesion (Martinez, 2013). Previous studies have noted the absence of flock formation in the absence of certain key species (Cody & Diamond, 1975; Maldonado-Coelho and Marini, 2000). Species that engage in high levels of information exchange with others can act as keystone species within multi-species flocks, influencing the behaviour and spatial dynamics of other members (Martinez, 2013). Therefore, prioritizing the conservation of these species is crucial for preserving the composition and functionality of avian communities (Martinez, 2013).

Although our study provides valuable insights into the formation of MSF and avian behaviour, it is important to recognise its limitations. Our observations were based on two years of observing flocks and two months of collecting acoustic samples, emphasizing the necessity for long-term data to strengthen the reliability of our findings. Future research should involve longer sampling periods and investigate the functions of specific vocalizations or the vocalizations of particular species, potentially through playback experiments or vocal muting experiments (McDonald 1989).

By incorporating measures of information flow into future studies on MSF formation and dynamics, we can gain valuable insights into the underlying mechanisms of avian social behaviour and contribute to more effective conservation strategies aimed at preserving biodiversity and ecosystem stability. In summary, our study adds to the expanding knowledge of MSF formation and underscores the importance of key species in driving flock dynamics.

Understanding the complex interactions among individuals and species within different habitats is critical for the effective conservation and maintenance of community structure. Further exploration of the vocalizations of MSF birds and their influence on flock behaviour holds the promise of unravelling the intricacies of this fascinating ecological phenomenon, paving the way for more comprehensive conservation strategies.

## ACKNOWLEDGEMENTS

We extend our heartfelt thanks to Pranav Balasubramanian, Anisha Mandal, and Tarun Menon for their unwavering support during the fieldwork period. Our gratitude also goes to the Arunachal Pradesh Forest Department for granting permission for this study. In addition, we sincerely appreciate the support from our peers, family, and the larger research community.

## AUTHORS CONTRIBUTION

**Satyam Gupta:** Conceptualization, Methodology, Formal Analysis, Investigation, Data Curation, Writing – Original Draft, Writing – Review & Editing, Visualization, Project Administration. **Akshay Bharadwaj:** Conceptualization, Methodology, Formal Analysis, Investigation, Writing – Review & Editing, Project Administration. **Umesh Srinivasan:** Conceptualization, Methodology, Writing – Review & Editing, Project Administration, and Supervision. **Akshiti Bhat:** Data Curation**. Aiti Thapa:** Data Curation. **Aman Biswakarma:** Data Curation. **Bharat Tamang:** Data Curation. **Binod Munda:** Data Curation. **Biren Biswakarma:** Data Curation. **Dema Tamang:** Data Curation. **Dambar Pradhan:** Data Curation. **Mangal K Rai:** Data Curation. **Rohit Rai:** Data Curation. **Shambhu Rai:** Data Curation.

## CONFLICT OF INTEREST

The authors have declared that no competing interests exist.

## Supplementary Material

**Table 1.**
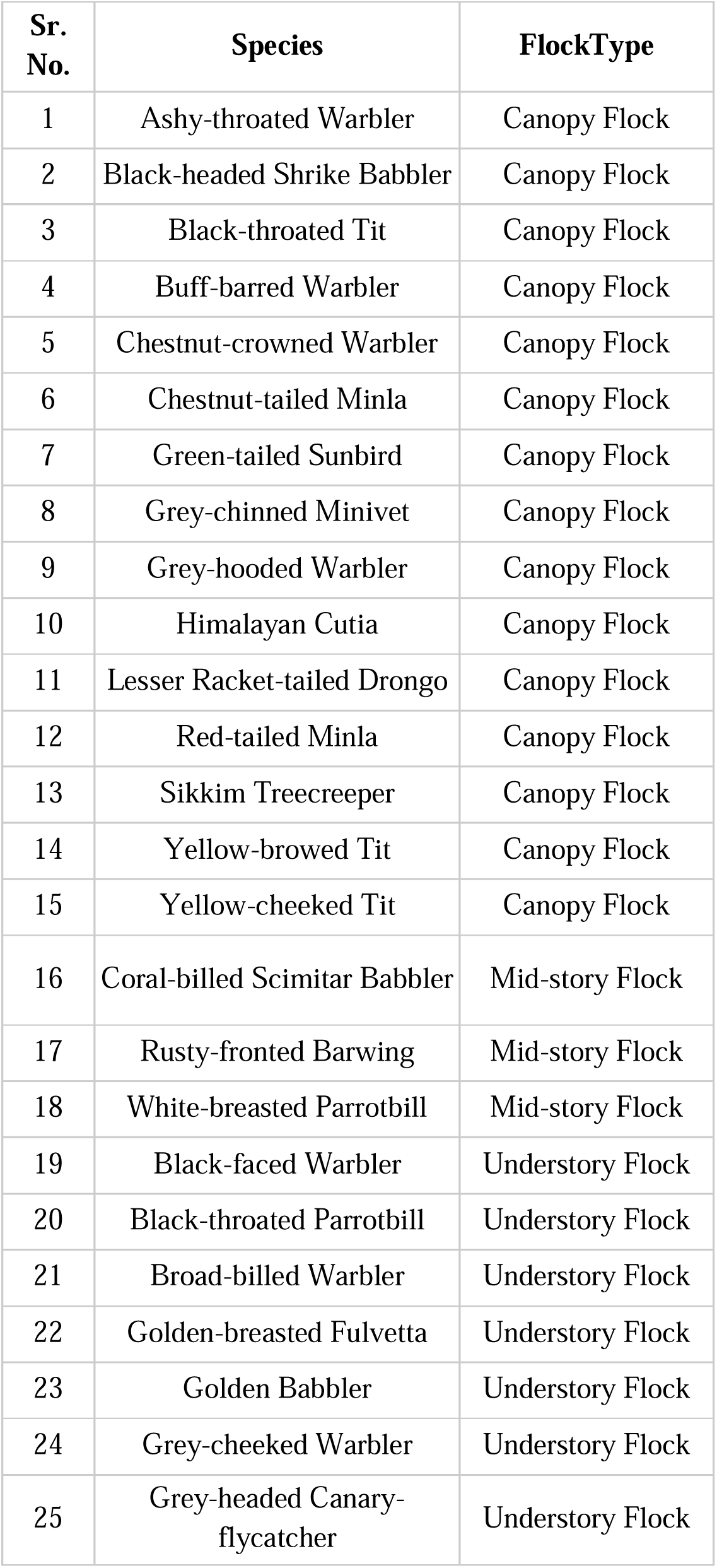
MSF participating birds in different flock types

**Figure 1.**
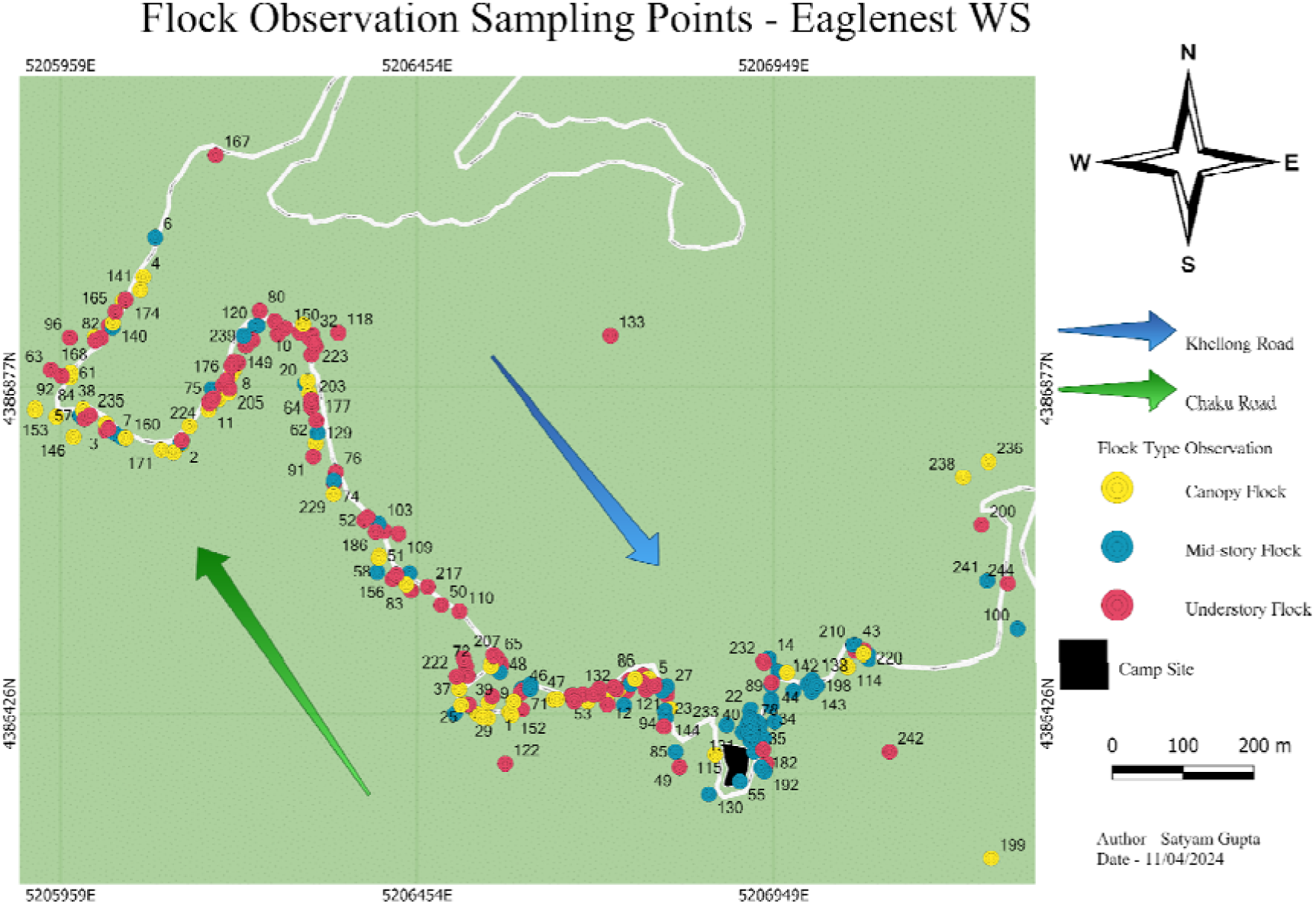
MSF Observation Sampling Locations

**Figure 2.**
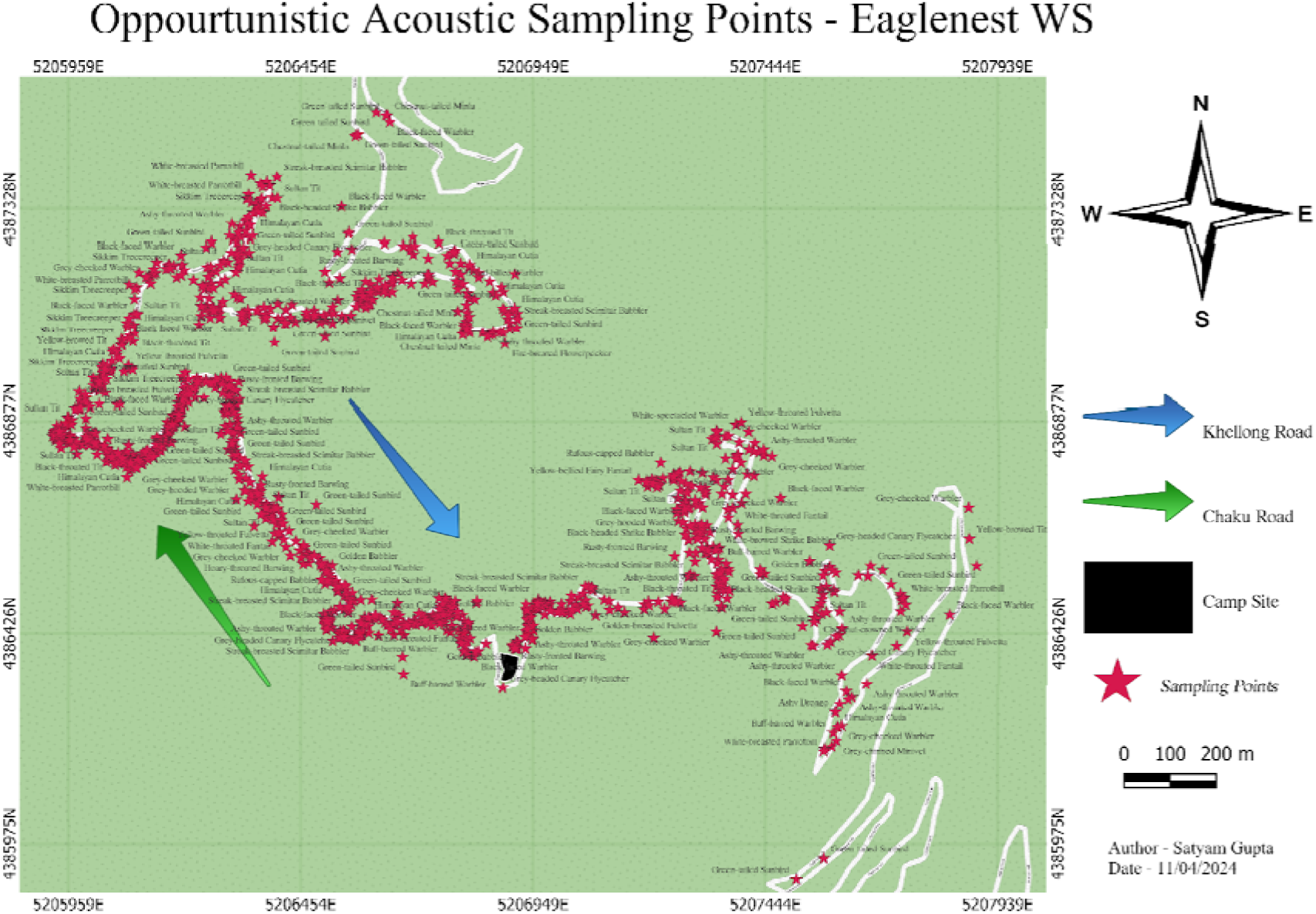
Opportunistic Acoustic Sampling Locations

**Figure 3.**
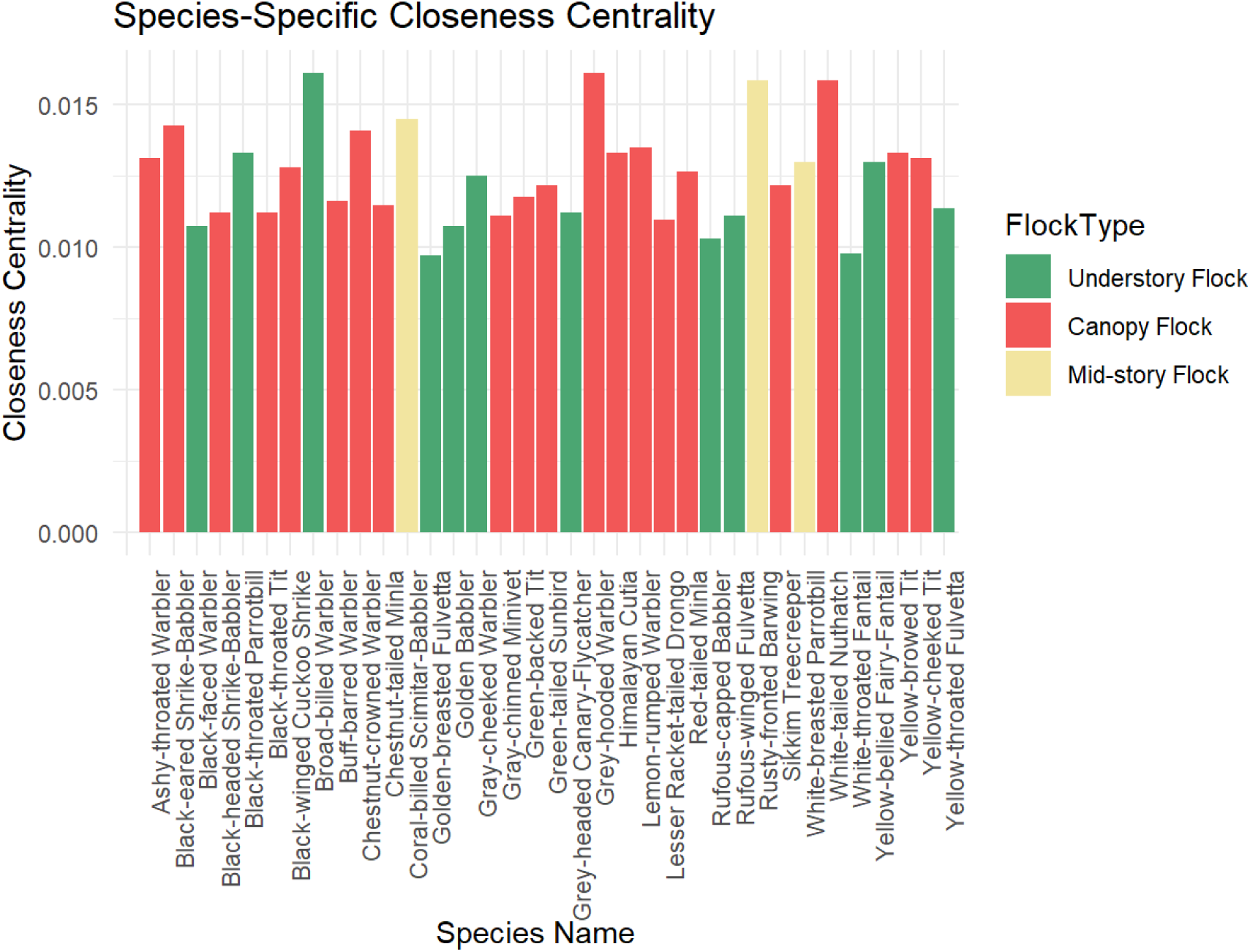
Distribution of Closeness Centrality in the MSF

**Figure 4.**
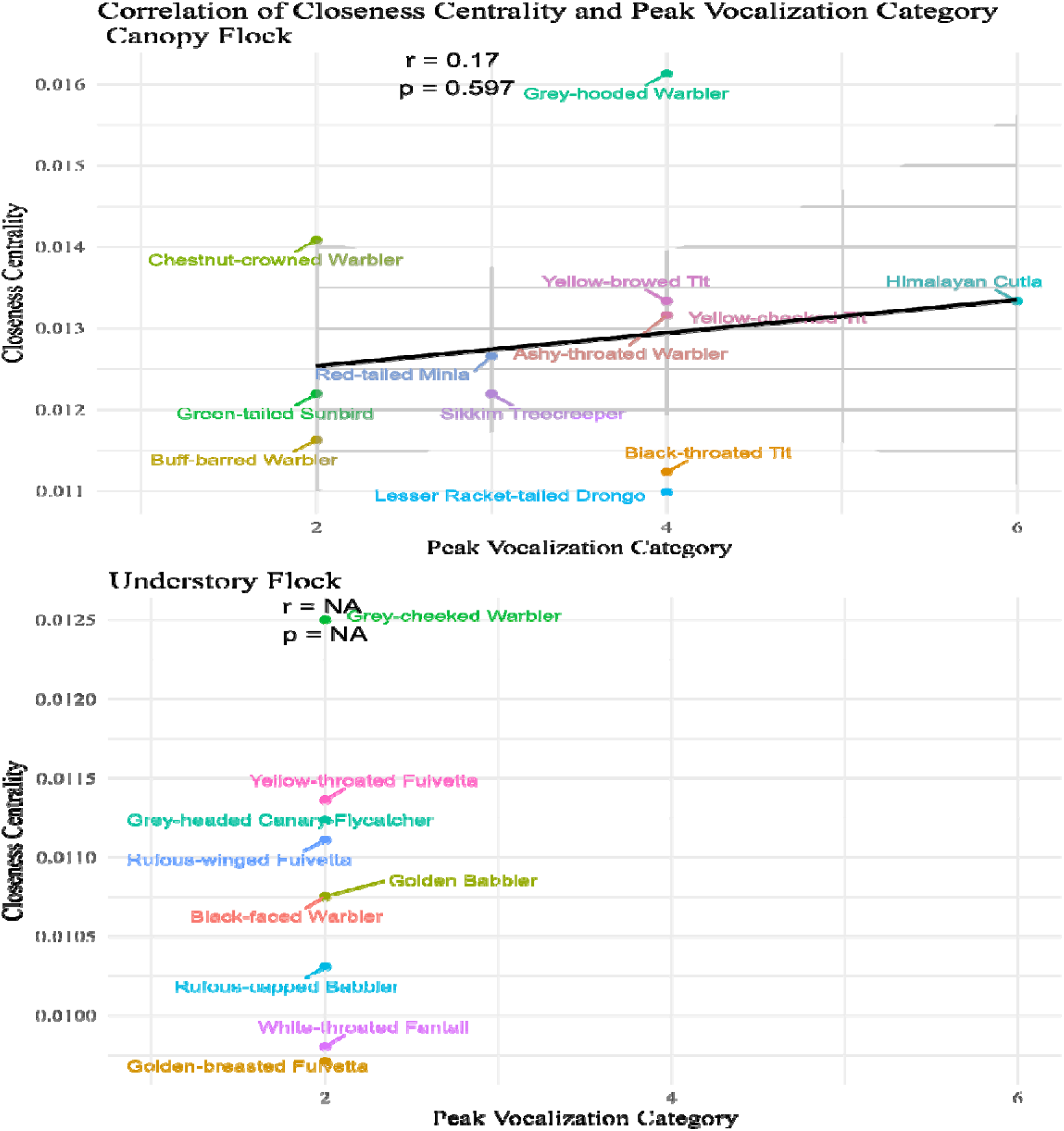
Correlation of Peak Vocalisation Period and Closeness Centrality. The Y-axis illustrates the values of closeness centrality, whereas the X-axis displays the peak vocalisation period. The plots indicate a positive correlation between closeness centrality and peak vocalization period for canopy flocks, although this relationship was insignificant. A correlation could not be established for the understory flock, as all variables on the X-axis were identical, indicating a synchronised peak vocalisation between the different species.

## Notes

### Competing Interest Statement

The authors have declared no competing interest.

### Summary of Updates

Akshay Bharadwaj's ans Satyam Gupta's affiliation was corrected.

